# Structure prediction of protein-ligand complexes from sequence information with Umol

**DOI:** 10.1101/2023.11.03.565471

**Authors:** Patrick Bryant, Atharva Kelkar, Andrea Guljas, Cecilia Clementi, Frank Noé

**Affiliations:** Department of Mathematics and Computer Science, Freie Universität Berlin, Arnimallee 12, 14195 Berlin, Germany; Department of Physics, Freie Universität Berlin, Arnimallee 12, 14195 Berlin, Germany; Microsoft Research AI4Science, Karl-Liebknecht Str. 32, 10178 Berlin, Germany

## Abstract

Protein-ligand docking is an established tool in drug discovery and development to narrow down potential therapeutics for experimental testing. However, a high-quality protein structure is required and often the protein is treated as fully or partially rigid. Here we develop an AI system that can predict the fully flexible all-atom structure of protein-ligand complexes directly, given a multiple sequence alignment representation of the protein and a SMILES string representing the ligand. At a high accuracy threshold, unseen protein-ligand complexes can be predicted more accurately than for RoseTTAFold-AA, and at medium accuracy even classical docking methods that use known protein structures as input are surpassed. The high accuracy presented here suggests that the goal of AI-based drug discovery is one step closer, but there is still a way to go to fully grasp the complexity of protein-ligand interactions. Umol is available at: https://github.com/patrickbryant1/Umol

## Introduction

Docking of small molecules to protein targets is an important problem for the evaluation of new drugs and the repositioning of known ones [1]. However, existing docking methods have significant limitations: (i) A high-quality structure of the protein is needed as the protein is usually treated at least partially rigid. (ii) The problem of identifying the correct docking pose is not solved [2]. (iii) Most evaluations are performed on structures in their bound (holo) form, limiting the search for new ligands to those which have identical binding modes to known ones [3]. A system that could predict the entire protein-ligand complex structure from a given protein sequence and the chemical structure of the ligand would address these challenges.

Recently, machine learning has been applied to the problem of protein-ligand docking [2,4–6]. However, these systems have not yet outperformed classical methods based on scoring functions [7–9] when considering a known target area or “pocket” [10]. This is a relevant test case as designing new drugs involves targeting specific binding sites on proteins [11], hence one can assume that the binding pocket is known.

On the other hand, it is not reasonable to assume that a protein structure is available in the bound (holo) form consistent with the ligand. When considering structures predicted with ESMfold [12], the success rate (SR, ligand ≤RMSD 2 Å) decreases to half of that on holo-structures (20.3% vs 38.2% of structures) using current state-of-the-art methods [2]. This suggests that pure protein structure prediction tools are not able to produce structures that are suitable for ligand docking.

Evaluation sets partitioned on release date and not on structural similarity is another confounder. When considering receptors dissimilar from those seen during training the SR is about half of that on seen holo receptors (20.8%) [2]. Considering unseen structures and the chemical validity (bond lengths and angles) of the ligands, the SR of some methods can go from 51% to only 1% [13]. Evaluating the same methods on unseen apo (unbound) structures likely results in even lower performance.

Protein flexibility is crucial to be able to reach the holo state and for successful ligand docking. Recently, an all-atom version of RoseTTAFold has been developed. RoseTTAFold All-Atom (RFAA) [14] allows for predicting proteins in combination with ligands and other biomolecules. However, the SR on the PoseBusters’ test set [13] for protein-ligand prediction is 42%, and drops to 23% for proteins that have not been seen during training (sequence identity <30%) [14]. This suggests that the challenge of protein-ligand prediction is not yet solved.

## Results

Here we develop a protein-ligand co-folding network as a first step towards a **U**niversal **mol**ecular framework, **Umol** (Figure 1a). Starting from a protein sequence, protein target positions (pocket) and ligand SMILES, a multiple sequence alignment (MSA) and a bond matrix are created. From these, features are generated within the network and a 3D structure is produced. There are no limitations on the flexibility of either protein or ligand since no structural information is required to produce the final protein-ligand complex structure.

**Figure 1.**
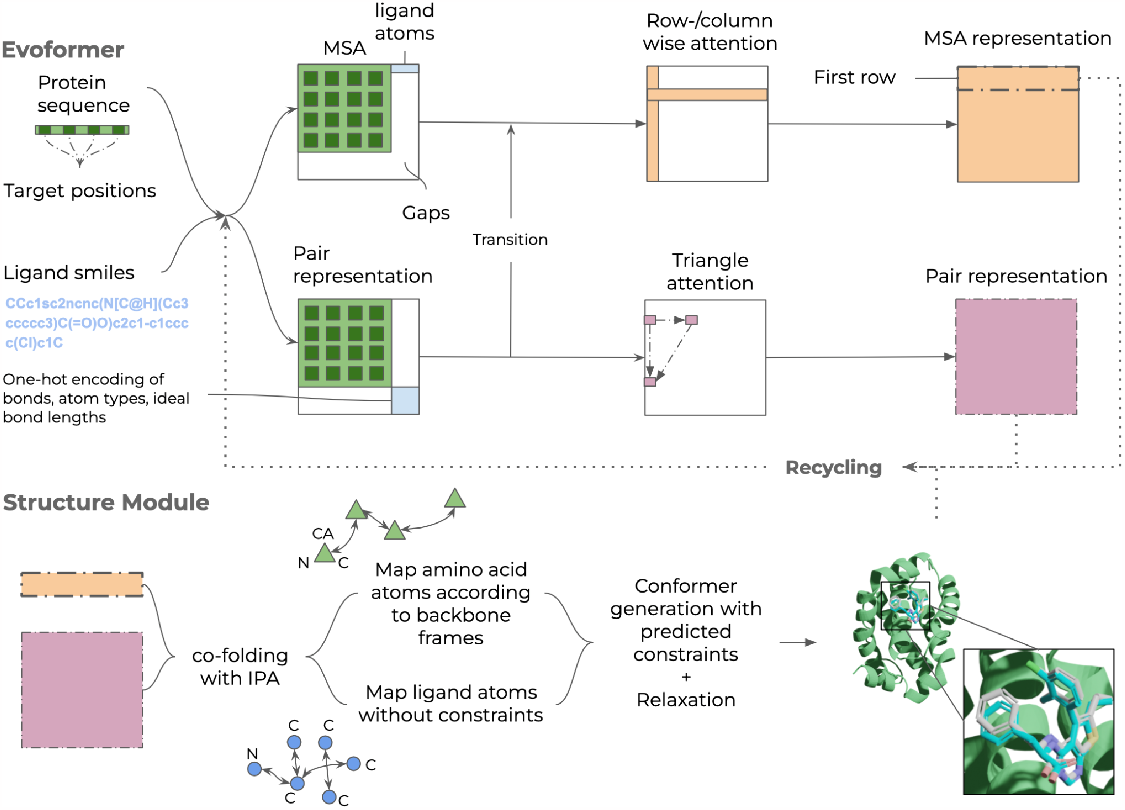
Description of Umol. An Evoformer network that processes both protein and ligand atom information. There are 48 Evoformer blocks. The protein is represented by a multiple sequence alignment (MSA) and target positions in the protein (pocket, Cβs within 10 Å from the ligand) are also defined. The ligand is represented by a SMILES string. Two different tracks are present. The top track processes the MSA (gaps for the ligand) and the bottom track processes pairwise connections within and between the protein and ligand. The resulting representations are fed into the structure module (8 blocks) which produces a 3D structure of the protein-ligand complex. The entire network is trained end-to-end and the representations and predicted atom positions are recycled to refine the final structure. An example for PDB ID 7NB4 (ligand RMSD=0.57) is shown with the predicted protein in green, the predicted ligand in blue and the native ligand in grey. The ligands are coloured by atomic types (blue=nitrogen, red=oxygen, remaining=carbon).

### Protein-ligand structure prediction

Figure 2a shows the success rate (SR, the fraction of predictions with a Ligand RMSD≤2 Å [15,16]), on 428 diverse protein-ligand complexes [13] for eight protein-ligand docking methods in addition to Umol. Umol and RoseTTAFold All-Atom (RFAA) [14] are the only methods that do not require the native protein structures as input and achieve 45.3% and 42% SR, respectively. The best method is AutoDock Vina [7] with 52.3% SR but it requires the native protein structure as an input.

**Figure 2.**
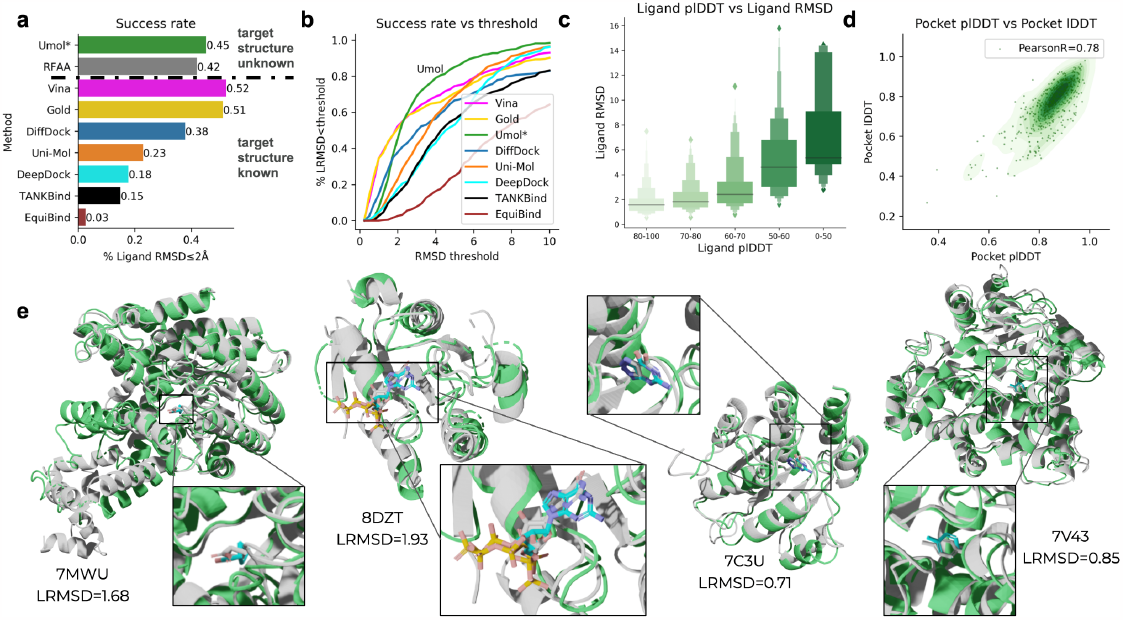
**a)** Success rates (fraction of predictions with ligand RMSD≤2Å) on the PoseBuster benchmark set. Note that only Umol (the method developed here) and RoseTTAFold All-Atom (RFAA) do not require the native protein structures as input (target structure unknown). **b)** Success rate vs ligand RMSD threshold. Umol has many complexes right above the 2 Å mark, suggesting that many ligands are almost in their native configuration. At a threshold of 3Å, the SR is 69% for Umol vs 58% for Vina. **c)** Ligand plDDT vs ligand RMSD for structures predicted with Umol. The success rate (ligand RMSD≤2 Å) for each bin is 80-100: 72.3%, 70-80: 58.2%, 60-70: 34.4%, 50-60: 5.3%, 0-50: 0.0%. **d)** Protein pocket predicted lDDT (plDDT) vs pocket lDDT for structures predicted with Umol as a density plot with each datapoint represented as a green scatter. The Pearson correlation coefficient is 0.78, suggesting a strong relationship between the two. **e)** Examples of predictions from the Posebusters test set with low homology to the Umol training set. The native structures are in grey and the predicted protein/ligand in green/cyan. The ligands are coloured by atomic types (blue=nitrogen, orange=phosphorus, red=oxygen, remaining=carbon). PDB IDS from left to right: 7MWU, 8DZT, 7C3U and 7V43 with ligand RMSDs 1.68, 1.93, 0.71 and 0.85, respectively.

A success cutoff of 2 Å ligand RMSD (LRMSD) is arbitrary. Figure 2b shows the SR vs the ligand RMSD threshold. Umol has many complexes right above the 2 Å mark, suggesting that many ligands are almost in their native configuration. At just above 2 Å (2.35 Å), Umol surpasses all other methods and at a threshold of 3 Å, the SR is 69% for Umol vs 58% for Vina. Umol has no successful complexes below 0.5 Å, but Vina and Gold do. This is likely a consequence of these methods using the native structures as an input, resulting in errors of close to 0 Å which should not be possible in a realistic setting.

It is important to evaluate a method on dissimilar proteins as similar proteins are likely to bind similar ligands in similar poses. Therefore, we divide the proteins on 30% sequence identity [13] (Table 1) to assess the performance on targets dissimilar to PDBbind from 2020 [17]. For the classical methods (AutoDock Vina, Gold) the performance is stable across protein sequence similarities, but for some AI-based methods, the performance is substantially reduced (e.g. 14.8% vs 48.0% for DiffDock). For Umol, the performance is 35.2% on targets with <30% sequence identity outperforming all other AI-based methods. No scores are available for RFAA, although their analysis indicates that the corresponding fraction is 23% [14].

**Table 1.**
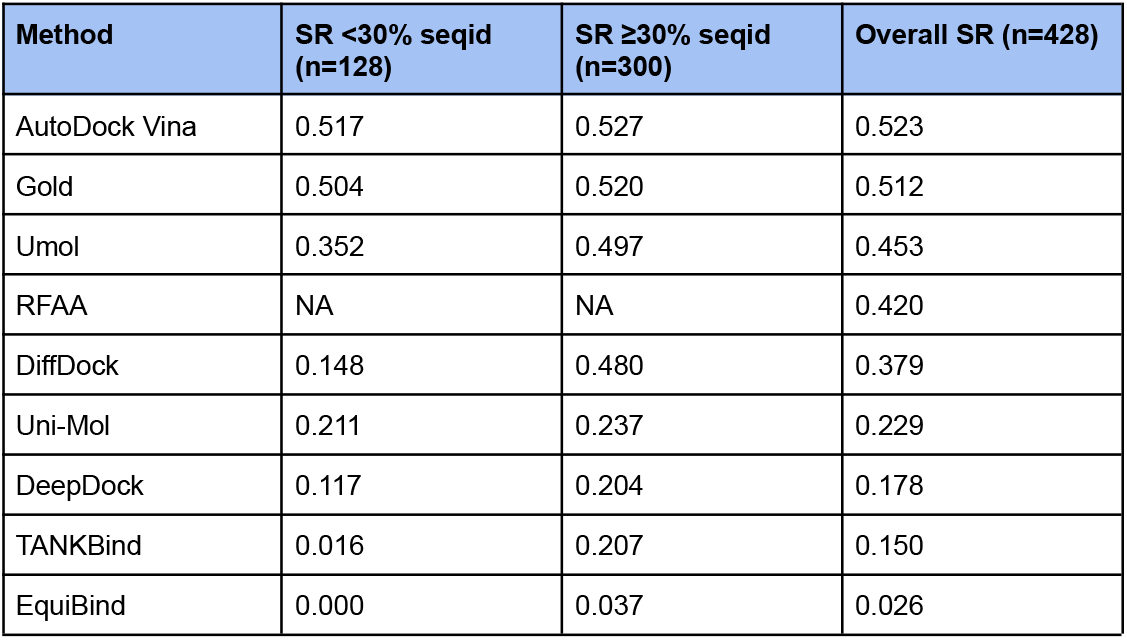
Success rate (% with ligand RMSD≤2Å) on the PoseBuster benchmark set divided by sequence identity (seqid) to the PDBBind 2020 release. The number of proteins with ≥30% seqid is 300 and <30% seqid 128 (n=428 in total). For Vina and Gold the performance remains high below 30% seqid. For diffdock, the performance is much lower below 30% seqid. Umol has the highest performance of the AI-based methods, also <30% seqid (35.2%). The estimated SR <30% seqid is 23% for RFAA. Note that RFAA and Umol are the only methods not receiving the native holo protein structures as input.

| Method | SR <30% seqid (n=128) | SR ≥30% seqid (n=300) | Overall SR (n=428) |
| --- | --- | --- | --- |
| AutoDock Vina | 0.517 | 0.527 | 0.523 |
| Gold | 0.504 | 0.520 | 0.512 |
| Umol | 0.352 | 0.497 | 0.453 |
| RFAA | NA | NA | 0.420 |
| DiffDock | 0.148 | 0.480 | 0.379 |
| Uni-Mol | 0.211 | 0.237 | 0.229 |
| DeepDock | 0.117 | 0.204 | 0.178 |
| TANKBind | 0.016 | 0.207 | 0.150 |
| EquiBind | 0.000 | 0.037 | 0.026 |

To see if accurate predictions can be distinguished from inaccurate ones based on the Umol model outputs, we analyse the relationship between its performance with the predicted lDDT [18] (plDDT, Figure 2c). At plDDT >80, the SR is 72% and <50 plDDT 0%, suggesting that accurate ligand poses can be distinguished. The same is true for the protein pocket plDDT displaying a Pearson correlation of 0.78 with the lDDT (Figure 2d). We also conclude that the proteins are predicted with high accuracy overall (Suppl. Fig. 1), reporting an average TM-score [19] of 0.96.

Fig 2e shows examples of predictions with low homology to the training set (<30% sequence identity) in structural superposition with the native complexes. The protein structure as a whole may differ in the tertiary structure (e.g. 7MWU, LRMSD=1.68), but Umol predicts the ligand in the correct binding pose. Some structures have flexible loops within the binding pockets (e.g. 8DZT, LRMSD=1.93), and Umol predicts these accurately as well. Other ligands bind within the protein structure (e.g. 7V43, LRMSD=0.85) and the protein structure is correctly predicted around the ligand.

### Difficult targets

To visualise what may distinguish complexes that are predicted well to those that are not, we select the bottom five complexes in terms of average LRMSD across all methods. We focus on targets with <30% sequence identity to PDBBind to adjust for overfitting. The bottom five complexes (highest LRMSD) are PDB IDs 8E77, 7QE4, 8F4J, 7MSR and 7JNB with Umol LRMSDs 11.74, 1.59, 13.31, 2.33 and 2.52 respectively (Figure 3). 7EQ4 is successful at 2 Å LRMSD and 7MSR and 7JNB are close suggesting that Umol can find correct poses even when other methods struggle. The larger, more complicated ligands 8W77 and 8F4J are predicted in the wrong orientations, suggesting that larger ligands are more difficult to dock. However, this relationship is not apparent when analysing all ligands (Supplementary Figure 2).

**Figure 3.**
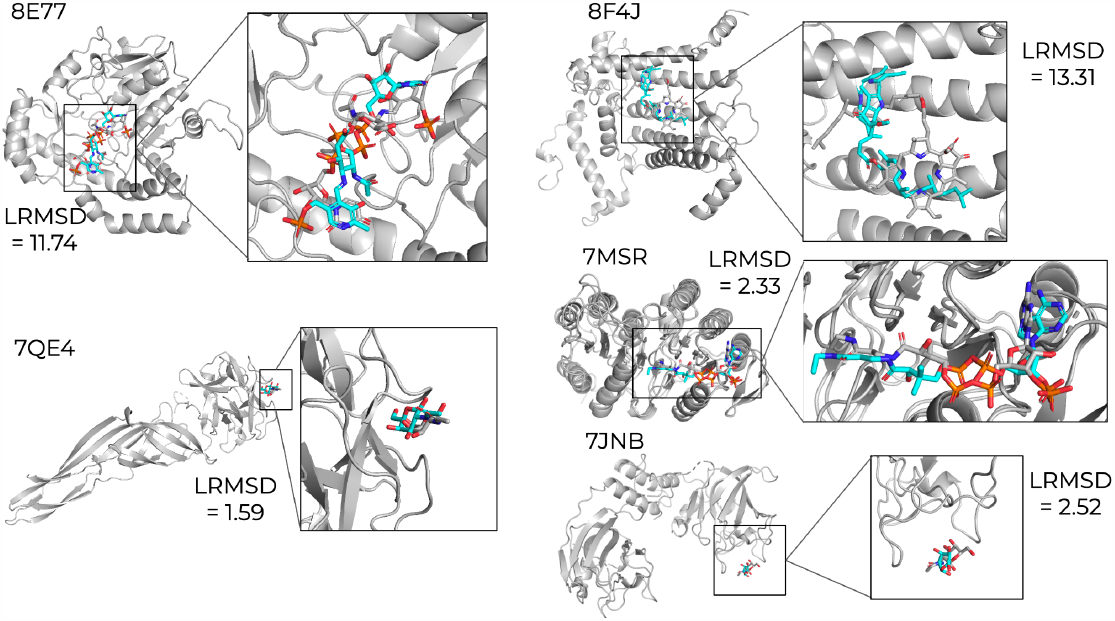
The 5 most difficult structures in terms of average LRMSD across all methods are 8E77, 7QE4, 8F4J, 7MSR and 7JNB with Umol ligand RMSDs 11.74, 1.59, 13.31, 2.33 and 2.52 respectively. The native structures are in grey and the predicted ligands in cyan.

## Discussion

We have introduced Umol, a neural network for predicting the complete all-atom structure of protein-ligand complexes only from protein sequence information, the position of the binding site in the sequence, and the chemical graph of the ligand. Umol does not depend on any structural information in contrast to all other ligand docking methods that rely on native protein structures or template information. Compared to its closest relative, RoseTTAFold All-Atom, Umol obtains a higher success rate (Ligand RMSD≤2 Å) on the PoseBusters test set (45% vs 42%) making it the highest performing method for protein-ligand structure prediction.

There are many poses slightly above the success threshold of 2 Å that are likely equivalent, which indicates that a more flexible scoring system may be required. This is exemplified by the fact that Umol surpasses the success rate of AutoDock Vina at a threshold of 2.35 Å. In cases where the native protein structures are not used for the scoring, even small errors in alignment become an issue.

Co-folding protein-ligand complexes has the potential to accelerate drug repositioning. In particular, we find that the predicted lDDT of the ligand can be used to select accurate docking poses and that the predicted IDDT of the protein pocket is suitable to select accurate interfaces (Figure 2). By inverting the trained network, it is possible that new ligand-binding proteins or protein-binding ligands can be designed. Another option is using transfer learning to create a generative diffusion model for the same purposes [14,20].

We do not find clear relationships between the model’s predictive performance and different features related to the protein or ligand (Suppl. Fig. 2). However, we do find that among the cases where other methods struggle, Umol is accurate in 3 out of 5 cases (Figure 3). The cases among the 5 where the performance is bad for Umol is when the ligands are large and complicated, resulting in wrong orientations.

The current release of PDBbind contains data processed from PDB in 2019. Since then, many more protein-ligand complexes have been submitted suggesting that higher accuracy may be possible to achieve. However, it is unclear what accuracy is needed to obtain meaningful results for protein-ligand docking. It is evident that the high accuracy observed in protein structure prediction is not obtained for tasks involving other molecules such as small molecules or RNA [21–23]. Without coevolutionary information on proteins, the accuracy of the structure prediction decreases rapidly [12,24]. As there is no similar source of information for small molecules or RNA, one has to rely solely on atomistic representations.

There is still a long way to go to grasp the complexity of protein-ligand interactions fully, but leveraging deep learning for structure prediction of the entire complex may bring us one step closer to the solution.

## Methods

### PDBbind

We used PDBbind from 2019 (2020 release [17]) processed by the authors from EquiBind (https://zenodo.org/record/6408497, 19119 protein-ligand complexes). We parsed all protein sequences from the PDB files. 18884 out of 19119 protein structures (99%) could be parsed (<80% missing CAs and >50 residues). Only the first protein chain in all protein-ligand complexes used here and in the evaluation was extracted. Features (see below) could be generated for 17936/18884 (95%) protein-ligand complexes. The failed ones did so due to issues of converting SMILES to 3D structures using RDKit (version 2023.03.2, https://www.rdkit.org).

#### Data partitioning

We cluster all protein sequences with MMseqs2 (version f5f780acd64482cd59b46eae0a107f763cd17b4d) [25] at 20% sequence identity with the options:

~~~
mmseqs easy-cluster DB.fasta clusterRes tmp --min-seq-id 0.2 –c
0.8 --cov-mode 1
~~~

We obtain 1486 sequence clusters at 20% sequence identity, representing the number of possible protein folds. Most sequences are below 1000 residues (median=263 residues, Figure 4a) and most clusters have only a few entries (Figure 4b). We select 90% of the clusters for training, 5% for validation and 5% for eventual calibration tasks (Table 2).

**Table 2.** Data partitioning for training, validation and calibration.

| Partition | Number of clusters | Number of protein-ligand complexes |
| --- | --- | --- |
| Train | 1486 | 16420 |
| Valid | 82 | 845 |
| Calibration | 84 | 671 |
| Total | 1652 | 17936 |

**Figure 4.**
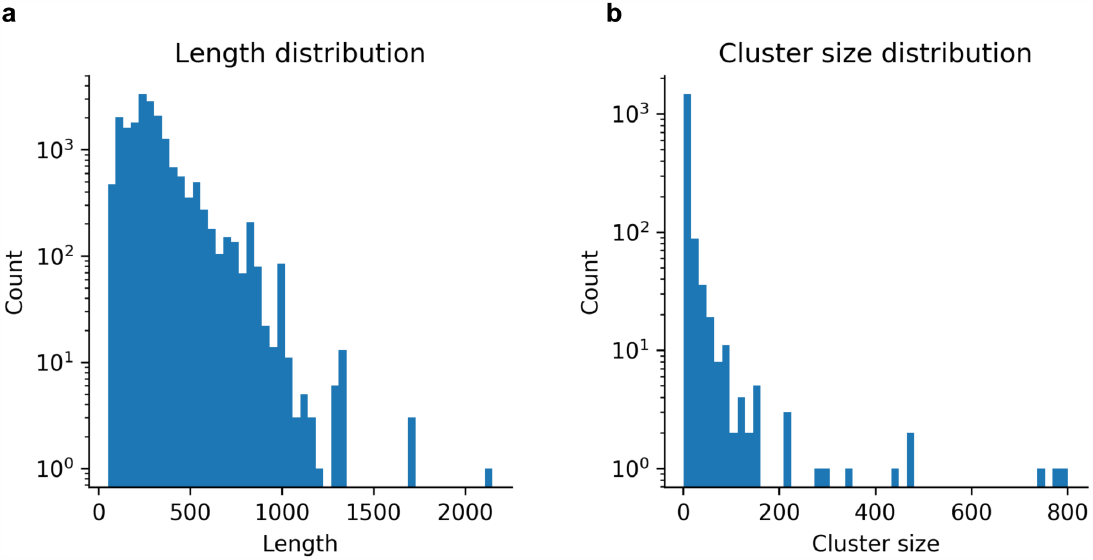
**a)** Length distribution (n=17936, median length=263 residues). **b)** Number of sequences in each 20% sequence identity cluster (median cluster size=2).

Note that the main evaluation and comparison with other methods(Figure 2, Table 1) is performed on the PoseBusters benchmark set that only includes structures not included in PDBbind 2020 (described below). The partition of the data in PDBBind was made before the PoseBusters benchmark was available, whereupon a decision to use that test set instead was made.

### PoseBusters test set

We evaluate the trained network and compare it with 8 other methods for protein-ligand docking on the PoseBusters benchmark set which contains 428 complexes not present in the PDBbind 2020 release [17] used for training here. We also compared the performance by dividing the dataset based on the sequence overlap to the training set (Table 1) [13].

We downloaded the PoseBusters set and scores for all methods except for RFAA from https://zenodo.org/record/8278563. The success rate for RFAA was taken from their preprint [14].

15 complexes are out of memory using an NVIDIA A100 GPU with 40Gb of RAM during inference (above 1000 residues). We crop these to 500 residues to include as many of the target residues (pocket) as possible.

### Network Description and Inference

The network architecture is a modification of the EvoFormer used in AlphaFold2 [24]. For an overview of the network see Figure 1. Note that no template information is used in the network, making it purely sequence-based. Here we outline the encoding and processing of the different network inputs.

#### Pocket encoding

All CB (CA for Glycine) within 10 Å from any ligand atom are one hot encoded. This is an arbitrary threshold similar to that of DeepDock [26]. Likely, similar thresholds will perform equally well. The encoding is used to bias both the MSA and pair representations and is processed throughout the network.

#### Ligand encoding

All atoms present at least 100 times in the ligands of the training dataset (B, C, F, I, N, O, P, S, Br, Cl) are one hot encoded. All other atoms are encoded with the same one hot encoding representing rare ligand atoms (As, Co, Fe, Mg, Pt, Rh, Ru, Se, Si, Te, V, Zn). We decided not to encode rare atoms to not overfit to these and to save space as this would result in a much sparser atom encoding.

The ligand bonds are one hot encoded as well. We encode single, double, triple and aromatic bonds separately and all other bonds as rare.

#### MSA generation and processing

To generate multiple sequence alignments, we search uniclust30_2018_08 [27] with HHblits (from HH-suite [28] version 3.1.0) with the options:

~~~
hhblits -E 0.001 -all -oa3m -n 2
~~~

We add a gap for the ligand atoms in the MSA representation and process the MSA in the MSA track alone. The MSA track is aware of the ligand through interactions with the pair track and due to the size and encoding of this in the initial MSA representation.

#### Pair processing

We process the pair representation with the atoms directly: amino acids+atoms, and let the MSA information flow to the pairwise interactions to influence the folding.

#### Recycling operations

The pair representation, the first row of the MSA representation and intermediate predicted final atom positions are recycled through the network 1-3 times sampled uniformly during training. For the predictions, three recycles were used for the validation set.

#### Loss

We use the same main loss functions and loss weights as in AlphaFold2, but with slight modifications to better suit the problem of protein-ligand structure prediction. The loss used up to step 24500 is

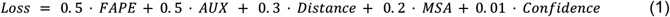

Where *FAPE* is the frame aligned point error, *AUX* a combination of the *FAPE* and angular losses, *Distance* a pairwise distance loss, *MSA* a loss over predicting masked out MSA positions and *Confidence* the difference between true and predicted lDDT scores. These losses are defined exactly as in AlphaFold2 and we refer to the description there [24].

The FAPE is calculated using only the amino acid N-CA-C frames towards ligand atoms as well. The MSA loss is only applied for the protein as well, but the distance and aux loss include both protein and ligand.

All ligand atoms are represented as their own frames (like a gas of atoms without constraints). The alternative is to enforce bond distances and known geometric properties. To enforce accurate bond lengths, we introduce an L2 distance loss at step 24500. This L2 distance loss is defined using the distance from the ideal ligand bond lengths extracted from a generated conformer in RDKit (clipped at 10 Å). The reason for not including this initially is to let the network learn freely how the protein and ligand should interact. We find that imposing too many constraints too early impedes the training and may lead to local minima according to initial tests (not shown). This loss has a weight of 0.1 and is added into the AUX loss. This loss acts as a form of harmonic potential for the ligand bonds.

#### Training and validation

We sample the sequences with inverse probability to the 20% sequence identity cluster size. We use a learning rate of 0.001 with 1000 steps of linear warmup and clip the gradients with a global norm of 0.1 as in AlphaFold2 [24]. The optimiser is Adam [29] applied through the Optax package in JAX (JAX version 0.3.24, https://github.com/deepmind/jax/tree/main). We train the network for 50000 steps (until convergence). Each step takes approximately 22 seconds, resulting in a total training time of 13 days. We validate every 10000 steps and assess the ligand RMSD and the protein pocket RMSD using the relaxation and scoring procedures described below.

The loss function declines rapidly (equation 1, Figure 5a). The masked MSA loss saturates quickly (Figure 5b), while the distogram loss is noisy throughout the training procedure (Figure 5c). The structure module loss (Figure 5d) is not saturated at step 50000, although the lDDT [18] for the protein and ligand only improves slowly (Figure 5e). Figure 5f suggests that the network starts to overfit to the training set between steps 40000-50000. The success rate on the validation set is the highest at step 40000 (26%).

**Figure 5.**
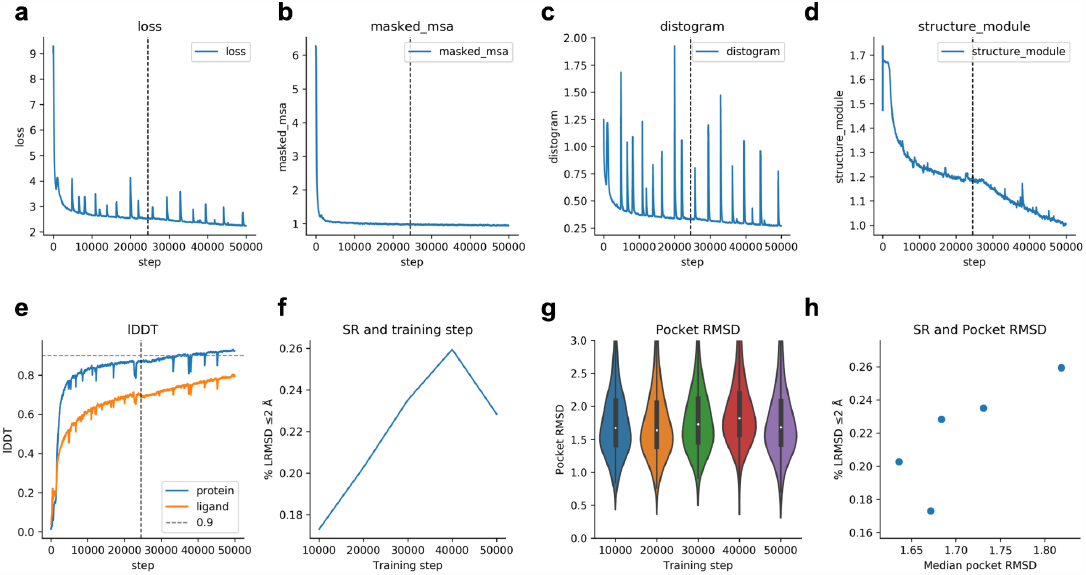
Training curves and metrics. The dashed vertical line indicates the time point where the additional L2 ligand distance loss was introduced (step 24500). **a)** The combined loss function (equation 1) vs the training step. **b)** Masked MSA loss vs training step. **c)** Distance loss vs training step. **d)** Structure module loss vs training step. This is the combination of FAPE and AUX (equation 1) **e)** lDDT for protein and ligand vs training step. The protein accuracy is higher than that of the ligand. **f)** Success rate (SR, % of protein-ligand complexes predicted with ligand RMSD≤2Å) for the validation set (n=741) at checkpoint intervals of 10000 steps. The examples that failed (n=104) did so due to RAM limitations and missing interface atoms. The SR is 17.3, 20.3, 23.5, 25.9 and 22.8 for steps 10000-50000, respectively. **g)** RMSD of the atoms in the protein pocket for the validation set (n=741) at checkpoint intervals of 10000 steps. The examples that failed (n=104) did so due to RAM limitations and missing interface atoms. **h)** Median pocket RMSD vs SR for the validation set (n=741) at the different checkpoints. The examples that failed (n=104) did so due to RAM limitations and missing interface atoms.

Both the lDDT of the protein and ligand are improving together, but there is a tradeoff in the SR. The pocket RMSD is the highest at the step where the SR is (Figure 5g and h), suggesting a tradeoff between these two. There is a difference of only 0.18 Å in the median pocket RMSD between the validation checkpoints, but the SR differs 8.6% (Figure 5h).

Conformer generation with RDKit is important to ensure that the predicted structures have realistic bond angles [13] and to adjust for small errors in e.g. the predicted protein side chain angles. Since the ligand atoms are represented as a point cloud, there are no constraints on bonds or angles resulting in possible violations in the predictions. To fix these issues, we align conformers generated by RDKit (version 2023.03.2, https://www.rdkit.org) from the input SMILES string using the ligand atom distance matrix as constraints to the predicted structures. We generate a total of 100 conformers and select the one with the lowest difference in atomic positions to the predicted positions.

### Relaxation with OpenMM

We noticed that some of the predictions contain clashes (here defined as two atoms being less than 1Å apart). This is a common problem with protein structure prediction [24], but can be easily amended using fast energy minimisation in a molecular dynamics force field, so-called relaxation. To relax the predicted structures, we add hydrogens to the protein and ligand and minimise the energy using OpenMM (version 8.0) [30]. We constrain the protein CA positions and the ligand positions, meaning that only the protein side chains are moved significantly.

The energy minimisation does not improve the ligand RMSD but fixes clashes and other errors within the protein. We use a Brownian Integrator and minimise the energy of the protein with a tolerance of 1 kJ for a maximum of 1000 steps with a restraining force of 100 kJ/nm^2^. This is a very fast relaxation procedure that essentially only alters side chain positions.

For the Posebuster test set, the relaxation resolves clashes in the protein-ligand interfaces. Before relaxation, 12% of interfaces have at least one clash, compared to 0.7% after the relaxation. The success rate is diminished by 2%, resulting in an SR of 43.3% compared to 45.3% before relaxation. This is a marginal change and we allow the user to determine if this is acceptable or not through the choice of relaxing the predicted structures or keeping them in their unrelaxed states.

### Timings

**Table 3.** Average time per process (the “real” time from the bash script is reported) needed for generating features and predicting the protein-ligand complex structures. The most time-consuming process is the MSA generation and the least is the network feature generation.

| What | Average time | Computational resources |
| --- | --- | --- |
| MSA generation | 285 s | 8 x 2.5 GHz Intel(R) Xeon(R) CPU E5-2680 v3 with 2.5 Gb RAM |
| Network input feature generation | 11 s | 2 x 2.5 GHz Intel(R) Xeon(R) CPU E5-2680 v3 with 2.5 Gb RAM |
| Protein-ligand structure prediction | 182 s | 1 x NVIDIA A100 GPU with 40 Gb RAM |
| RDKit ligand conformer generation | 13 s | 1 x 2.5 GHz Intel(R) Xeon(R) CPU E5-2680 v3 with 2.5 Gb RAM |
| OpenMM protein relaxation | 27 s | 1 x NVIDIA RTX A4000 with 16 Gb of RAM |
| Total | 518 s ≈ 9 minutes | At least 2 x 2.5 GHz Intel(R) Xeon(R) CPU E5-2680 v3 with 2.5 Gb RAM and 1 x NVIDIA A100 GPU with 40 Gb RAM |

### Comparison methods

We compare the performance on the PoseBusters benchmark [13] with the methods evaluated there for protein-ligand docking and the recently released RoseTTAFold All-Atom (RFAA). We describe the different methods briefly below:

#### AutoDock Vina

A classical docking method based on perturbing the ligand structure and calculating a score until convergence. Requires the native holo protein structure, a description of possible ligand poses, and a cube with a side length of 25 Å centred on the centre of mass of the heavy ligand atoms. [7]

#### Gold

A classical docking method based on perturbing the ligand structure and calculating a score until convergence. Requires the native holo protein structure, a description of possible ligand poses, and a cube with a radius of 25 Å centred on the centre of mass of the heavy ligand atoms. [31]

#### RoseTTAFold All-Atom

RoseTTAFold All-Atom (RFAA) is a neural network for predicting interactions between proteins and ligands as well as other atoms such as metal ions. RFAA lets each ligand atom move freely and treats the amino acids as N-CA-C frames as in Umol. The biggest differences to Umol are the “3D track”, the input of template and sterical information, the treatment of each ligand atom as an individual frame in the loss calculations (FAPE) and the lack of defining a pocket.

In Umol, the FAPE is only calculated as seen through the amino acid frames. This is because even though all ligand atoms can have frames, they will be highly variable depending on the predicted positions of the other ligand atoms. In contrast, the amino acid frames will be constant as the N-CA-C relationship remains the same regardless of the predicted positions.

As in Umol, RFAA uses relaxation of the predicted protein-ligand structures. RFAA uses Rosetta [32], while Umol uses molecular dynamics with OpenMM. Another difference in the loss calculations is that RFAA reorders all ligand atoms to find the lowest possible loss. The reason for not applying this here is that it is very difficult to exactly determine what atoms are interchangeable. Often, atoms can be interchangeable on a local scale, but not when considering the whole ligand. [14]

#### DiffDock

DiffDock takes ligand SMILES and the native holo protein structure as input and generates amino acid embeddings with ESM2. An initial ligand conformation is generated with RDKit which is updated and docked to the native protein structure using a diffusion process and deep learning. DiffDock does not require a defined pocket. [2]

#### Uni-Mol

A deep learning method that takes a ligand and native holo protein structure as input. Requires all protein residues within 6 Å of any heavy ligand atom. [33]

#### DeepDock

A deep learning method that takes a ligand and native holo protein structure as input. Requires a protein surface mesh of all protein residues within 10 Å of any heavy ligand atom. [26]

#### TankBind

A deep learning method that takes a ligand and native holo protein structure as input. Does not require a defined pocket. [4]

#### Equibind

An SE(3)-equivariant geometric deep learning method that takes a ligand and native holo protein structure as input. Does not require a defined pocket. [5]

### Scoring protein and ligand poses

To score the predicted protein-ligand complexes, we align the pocket CAs to the native structures and transform the predicted ligand accordingly. We then calculate the ligand RMSD and the RMSD of all available atoms in the protein pocket (equation 2). We assume that the atomic order is the same for the predicted/native ligands and do not adjust for symmetry. This can result in a lower success rate in some cases, but very few cases should be affected by symmetrical swaps beyond the 2 Å RMSD threshold.

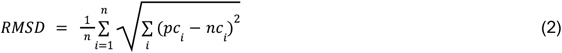

Where *pc* is the predicted 3D atomic coordinate (*x,y,z*) and *nc* is the native 3D atomic coordinate, and n is the number of heavy (non-hydrogen) atoms.

We score the overall protein structures with TM-align [34] and the command:

~~~
TMalign native.pdb predicted.pdb
~~~

### Number of effective sequences (Neff)

We clustered sequences at 62% identity to estimate the amount of information available in each MSA of the test set [35]. The resulting number of clusters were used as the number of effective sequences (Neff). We used MMseqs2 (version f5f780acd64482cd59b46eae0a107f763cd17b4d) [25] with the command:

~~~
mmseqs easy-cluster MSA clusterRes tmp --min-seq-id 0.62 -c 0.8
--cov-mode 1
~~~

## Acknowledgements

This study was supported by the European Commission (ERC CoG 772230 “ScaleCell”), MATH+ excellence cluster (AA1-6, AA1-10), Deutsche Forschungsgemeinschaft (SFB 1114 porjects A04 and C03, RTG 2433 Project Q04, SFB/TRR 186 Project A12), the BMBF (Berlin Institute for Learning and Data, BIFOLD) and the Einstein Foundation Berlin (Project 0420815101). Computational resources were obtained from ZIH (SCADS) at TU Dresden with project id p_scads_protein_na.

We thank the authors of PoseBusters for their extensive benchmark and for making the data available. We also thank the OpenMM team for discussions regarding the structure relaxation.

## Contributions

P.B designed and performed the studies, prepared the figures and wrote the initial draft of the manuscript. A.K performed the relaxation of the predicted structures in consultation with P.B and A.G. All authors contributed in reading and improving the manuscript draft. C.C and F.N obtained funding. P.B and F.N obtained computational resources.

## Availability

Umol is available at: https://github.com/patrickbryant1/Umol The predicted structures used for the calculation of various metrics presented in the figures here as well as the input features for the training of the network can be found at: https://zenodo.org/records/10060894

## Supplementary Material

**Supplementary Figure 1.**
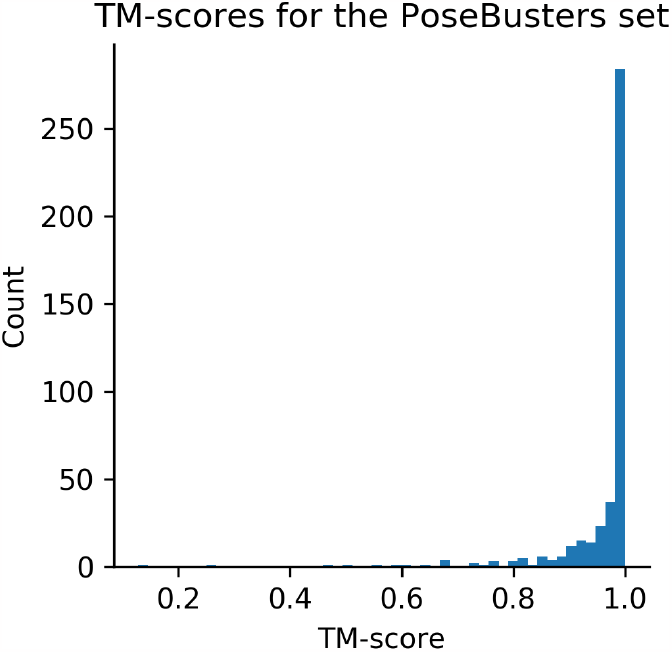
TM-scores for the PoseBusters test set comparing the predicted and native protein structures. The average TM-score is 0.96.

### What determines the ligand RMSD?

Targets with higher sequence identity to the training set tend to be predicted with higher accuracy (Table 1), but this does not fully explain the performance difference between targets. To see if certain features make the predictions more accurate, we analyse the conformer error when comparing the distance matrix of the ligand generated with RDKit (Methods) vs the predicted atomic positions, the ligand size, RMSD of the protein pocket atoms, the number of pocket residues, the number of effective sequences (Methods) and the protein length. We find only weak correlations among all features (**Supplementary Figure 2**), with the highest (Spearman R=0.56) being for the conformer error and the lowest for the protein length (Spearman R=0).

**Supplementary Figure 2.**
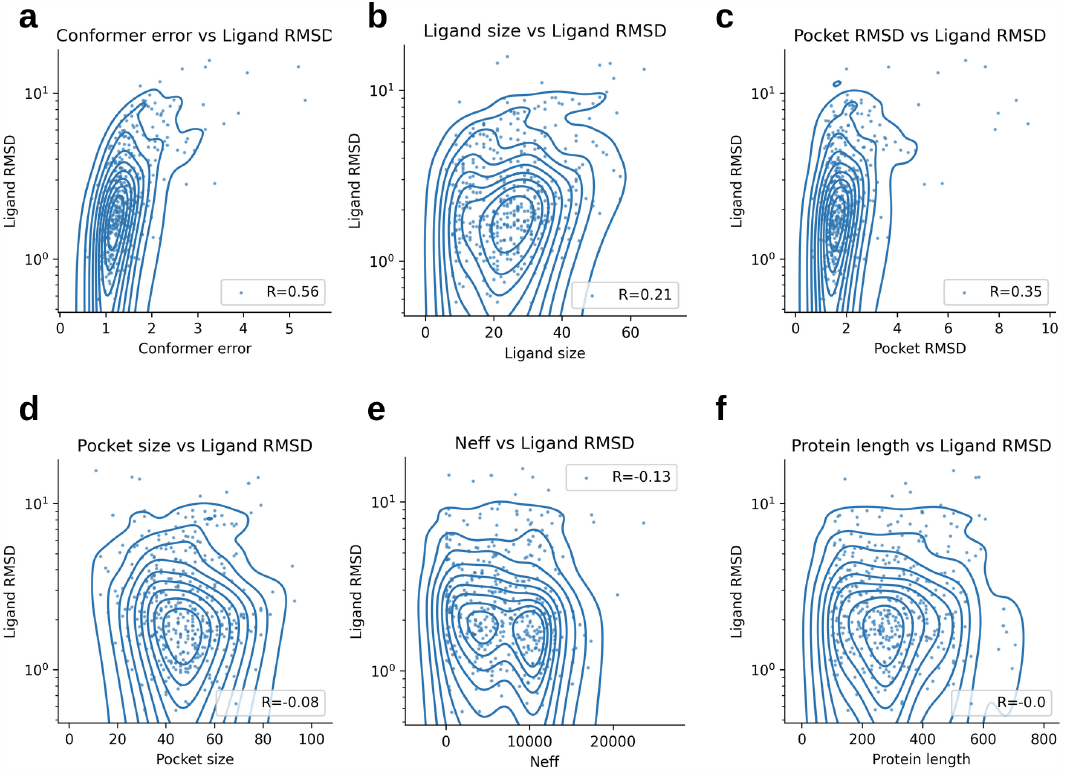
Analysis of the relationship between different features and ligand RMSD (LRMSD) for the predictions on the PoseBusters test set (n=428). The lines represent the density of the data points (dots). **a)** The error between the predicted constraints and the ligand conformer generated by RDKit (Conformer error) vs the LRMSD. **b)** Ligand size (number of atoms) vs LRMSD. **c)** RMSD of all the atoms in the protein pocket vs the LRMSD. **d)** The number of pocket residues vs the LRMSD. **e)** The number of effective sequences vs the LRMSD. **f)** The protein length (number of amino acids) vs the LRMSD.

## Notes

### Competing Interest Statement

The authors have declared no competing interest.

